# Effects of caller dominance rank on response to contact calls in ring-tailed lemurs (*Lemur catta*)

**DOI:** 10.64898/2026.10.11.758396

**Authors:** Nora Gerner, Roger Mundry, Oliver Gold, Kaja Wierucka

## Abstract

In group-living species, dominance hierarchies play a central role in shaping social dynamics, influencing both agonistic and affiliative interactions. Being able to recognise an individual’s dominance rank at a distance, can reduce direct conflicts, facilitating group coordination, and mediating social cohesion. While the role of dominance in relation to aggressive or reproductive vocalizations has been well documented, its influence on affiliative contact calls, particularly in mediating group movement and social integration, remains underexplored. We conducted playback experiments with free-ranging adult ringtailed lemurs (Lemur catta) at the Affenwald Wildlife Park to investigate how dominance rank of the caller and rank differences between caller and listener affect behavioral responses. Using Generalized Linear Mixed Models (GLMMs), we analysed three response variables: vocal response, number of visual scans, and occurrence of movement. Our results revealed sex-specific effects: females exhibited increased movement following contact calls from high-ranking individuals, and tended to refrain from moving when their own rank exceeded that of the caller. In contrast, no significant effects of dominance rank were observed in males. These findings suggest that high-ranking females may play an important role in initiating group movement. This study provides insights into the role of dominance rank in mediating social coordination through contact calls and provides a foundation for further research into the role of vocal communication in maintaining group cohesion in primates.

## Introduction

In social animals, dominance hierarchies serve as a key mechanism for organising interactions, minimising costly conflicts, and promoting social stability. Once established, these hierarchies allow individuals to adjust their behavior based on the rank of others, reducing the need for repeated aggressive encounters and thereby conserving energy and minimising injury risk (Schjelderup-Ebbe, 1922; Forkman & Haskell, 2004; Tibbetts et al., 2022). The ability to accurately assess one’s own rank and that of others is therefore a critical social skill, particularly in species with steep or stable dominance gradients. In such systems, individuals must continuously monitor and interpret social signals to navigate complex social landscapes, balancing cooperation with competition across a range of contexts – from access to food and mates to participation in collective decisions. While dominance information is often conveyed through visual or physical displays, acoustic signals are particularly advantageous due to their rapid transmission, long-range efficacy, and capacity to convey immediate social and physiological states (e.g., arousal, identity, rank). Indeed, vocalization structure or rate of their production have been shown to be linked to dominance rank (e.g., Sun et al., 2021; Bolt, 2021; Vannoni & McElligott 2008; Nogueria et al. 2016; Mathevon et al. 2010).

Research has primarily focused on the role of dominance in agonistic and reproductive communication, yet less is known about how rank influences responses to affiliative signals. Contact calls are used during routine social interactions and may support group cohesion and coordinated movement. Listeners may respond differently depending on the caller’s rank, whether rank is encoded directly in the call or inferred from the caller’s identity. For example, calls from high-ranking individuals may elicit greater attention or a stronger behavioral response, allowing these individuals to exert greater influence on collective decisions. In white-faced capuchins (*Cebus capucinus*), individuals respond more strongly to long-distance “lost calls” from dominant group members than to calls from lower-ranking individuals, allowing high-ranking individuals to rejoin the group more quickly and potentially reducing the risks associated with isolation (Digweed et al., 2007). Rank may also influence collective movement more generally. In Alaotran gentle lemurs (*Hapalemur alaotrensis*), higher-ranking individuals initiate group movements more often than lower-ranking individuals (Waeber and Hemelrijk, 2003). Understanding how social rank shapes responses to affiliative signals is therefore important for elucidating how animals coordinate collective behavior, maintain group cohesion, and navigate social relationships.

Here, we investigate the influence of dominance rank on social communication in ring-tailed lemurs (*Lemur catta*), a highly social and vocally complex primate species with a well-documented, female-dominated dominance hierarchy (Jolly, 1966; Kappeler, 1990). In this species, adult females consistently outrank all adult males. Ring-tailed lemurs possess one of the most extensive vocal repertoires among lemurs, including a wide array of contact calls used in everyday social interactions (Kittler et al., 2015). While previous research has demonstrated that dominance rank influences vocal behaviour in agonistic contexts, such as the increased production of “yip” calls by lower-ranking males during losing encounters (Bolt, 2021), the role of dominance in *affiliative* communication remains largely unexplored. Given the species’ pronounced female dominance, ring-tailed lemurs provide a valuable model for investigating whether listeners adjust their responses to affiliative calls according to the signaller’s rank, and whether this effect differs between females and males.

Affiliative contact calls in ring-tailed lemurs are critical for maintaining group cohesion, facilitating long-distance communication and coordinating group movement. These calls are often referred to as “cohesion calls” because they help individuals locate one another, maintain proximity, and reinforce social bonds—even when physical contact or grooming is not possible (Kulahci et al., 2015; Bolt & Tennenhouse, 2017). During group movement, contact calls are frequently produced by one individual, followed by vocal responses from others, which may then trigger synchronised movement (Macedonia, 1993; Oda, 1996). However, the precise mechanisms underlying movement initiation remain unclear. It is unknown which individuals are most likely to initiate calls, what determines whether a call leads to movement, or whether acoustic cues (e.g., the moan itself), visual cues (e.g., the initial movement of the caller), or a combination of both are necessary to prompt a response. Notably, moan exchanges sometimes occur without subsequent movement, suggesting a dual function: maintaining acoustic contact and signalling location, even in the absence of immediate group movement (Kondo & Watanabe, 2009). This functional ambiguity underscores the need to examine how social rank might modulate the *effectiveness* of such calls in eliciting behavioural responses.

In this study, we used playback experiments to test whether the dominance rank of the caller (both absolute rank and relative rank difference between caller and listener) influences the behavioural response of free-ranging ring-tailed lemurs. We hypothesised that high-ranking individuals would elicit stronger or more immediate responses, especially in movement-related behaviours, suggesting a leadership role in group coordination. Our findings provide novel insights into how dominance rank may be embedded in affiliative vocalisations.

## Methods

### Study site and population

Data were collected in a population of semi-free-ranging ring-tailed lemurs (*Lemur catta*) housed at the Affenwald Straußberg, Germany. The park houses about 80 (two groups of 35 to 40 individuals each) ring-tailed lemurs in a large, 3.5 ha outdoor enclosure that allows the animals to exhibit natural behaviours. The lemurs forage in the enclosure and are additionally provisioned with fresh food. Each individual within the park is equipped with a microchip, enabling identification. The animals are habituated to human presence. Only individuals from one of the groups were used in the study (Supplementary Materials, Table S1).

### Dominance ranks

To assess social dominance within the group, we conducted a series of controlled dyadic interaction tests using a modified food competition paradigm. 40 individuals were tested in June and July 2024, in about half of the possible unique dyadic combinations (N = 295), with each dyad tested only once. Prior to each test, two individuals who had not previously been tested together were isolated from the rest of the group, maintaining a minimum distance of approximately one lemur body length from non-participating individuals. The test was conducted in a neutral area, with the experimenter positioned approximately one meter in front of the dyad, holding a pair of tweezers with one raisin in each hand, spaced about one meter apart.

Once both individuals were visually attentive to the raisins, the experimenter slowly moved the two raisins toward the center over a 3-second interval, held them at the midpoint for 5 seconds, and then ended the trial. This standardised procedure ensured consistent stimulus presentation and minimized external interference. The outcome of each interaction was classified as follows: When only one individual accessed the food, that individual was classified as the winner and its partner as the loser. When food access did not provide a clear outcome, the individual that chased or physically displaced its partner through biting, slapping, or pushing (as defined by Pereira and Kappeler 1997) was classified as the winner. If one individual retreated while the other remained or approached, the retreating individual was classified as the loser. All interactions were video-recorded to ensure consistent outcome assignment.

Trials were excluded if a third individual approached within one lemur body length, if a raisin was taken before the end of the test, or if a dyad was tested more than once. To minimize behavioural carryover, no individual was tested twice in succession; instead, at least one intervening dyad or a 10-minute interval was enforced between consecutive tests involving the same individual.

The resulting dominance data were compiled into a binary matrix, with wins coded as 1, losses as 0, and ties as 0.5. To derive a ranked dominance hierarchy, we applied the Iterative and Simultaneous Optimization (I&SI) method (De Vries, 1998), implemented via the *ISI* function in the *Elo-Rating* package (ver 0.46.18; Neumann and Kulik, 2024) in R (R Core Team, 2024). Because the I&SI method assumes a linear dominance hierarchy, we first assessed the degree of transitivity in the dominance network using the triangle transitivity index (Shizuka & McDonald, 2012). Given that not all possible dyads were tested, resulting in a limited sample size, we ran the analyses 10 times with different random starting orders to assess stability. The final dominance rank for each individual was calculated as the average rank across the 10 runs. In order to obtain high values for high-ranking individuals and low values for low-ranking individuals, we subtracted the I&SI averaged value of all individuals from the maximum I&SI value in this study.

### Playback experiments

We used contact calls of ring-tailed lemurs (*Lemur catta*) characterized in the literature by a fundamental frequency with a median range of 336Hz to 480Hz and a total range of 256 to 960Hz, commonly referred to as “moans” (Macedonia, 1993). These vocalizations have also been described as “meows,” “wails,” or “hms” in prior studies (Andrew, 1963; Jolly, 1966; Oda, 1996; Bolt & Tennenhouse, 2017; Figure 1). The calls were recorded from adult individuals (4 to 15 years old; 8 females, 13 males) spanning a range of dominance ranks within the study group. Recordings were collected opportunistically during natural social interactions using a handheld omnidirectional microphone (AudioMoth 1.2.0 development board, AudioMoth, UK).

**Fig 1.**
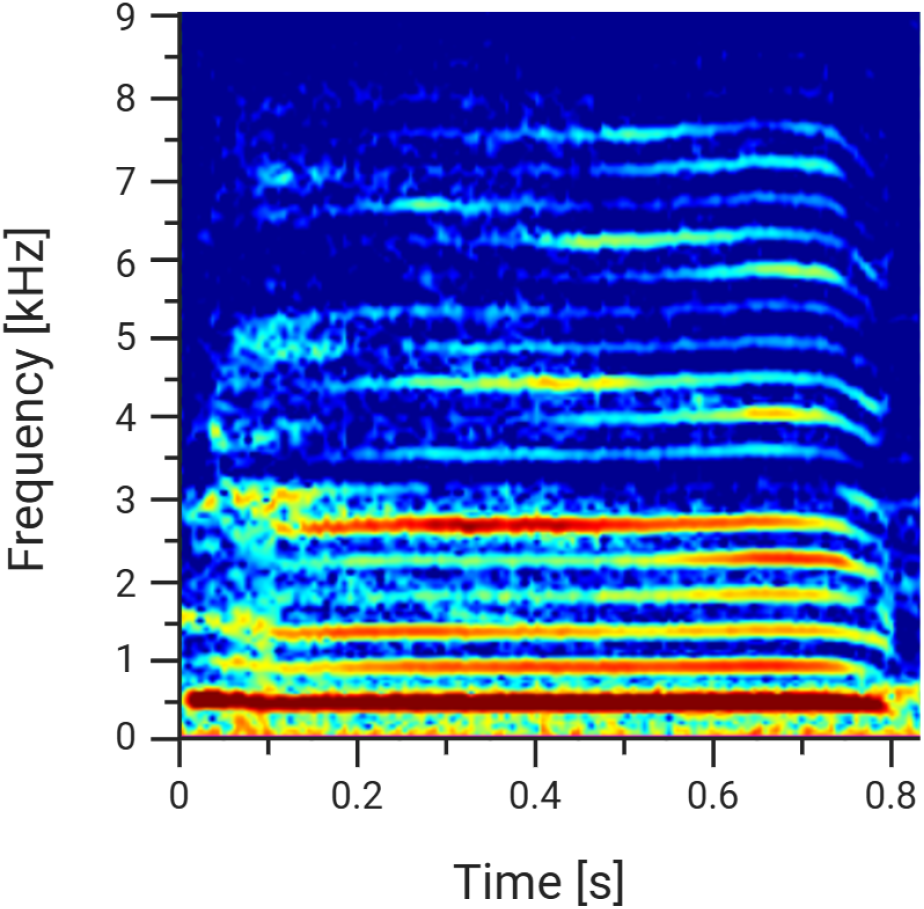
Example spectrogram of a ring-tailed lemur contact call used as playback call. Spectrogram settings: Hann window size of 1300 samples, hop size of 650 (overlap 50%).

To extract calls for playback recordings, we used Raven Lite 2.0.5 (K. Lisa Yang Center for Conservation Bioacoustics, Cornell Lab of Ornithology, 2024) for visual inspection and manual selection of high-quality vocalisations. Calls were excluded if they contained significant background noise or overlapping vocalisations. Selected calls were normalised to a peak amplitude of 1.0 dB using Audacity 3.7.4 (Audacity Development Team, 2025) to ensure consistent acoustic input across trials. Each playback stimulus consisted of two different calls from the same individual, separated by 5 seconds of silence. A total of 21 distinct playback stimuli were created, representing individuals of varying sex and dominance rank. These stimuli were used to test responses across a range of social contexts.

Playback experiments were conducted between July and September 2025. Each of the 18 focal individuals (aged 4 years) was exposed to 20 to 21 different playback calls, with no individual being played back their own vocalisation. Stimuli were presented in randomized order, and two experiments with the same animal were at least 1 hour apart. During each trial, the focal individual was isolated from the group at a minimum distance of 10 meters from all other group members. A speaker (Apple Beats Pill A1680) was positioned 1.5 meters from the focal animal, placed out of direct visual range. All trials were recorded using a GoPro HERO9 Black camera for later behavioural analysis.

Experiments were excluded from analysis if (1) a vocalization occurred within 10 seconds before playback onset; (2) another individual responded to the playback and the focal individual vocalised shortly afterward, making the source of the response ambiguous; (3) the focal individual was not visible on video (precluding reliable scoring of visual scans or movement); (4) the experiment was disrupted by other lemurs, park visitors (within the 10-meter radius); or (5) the focal individual approached, or touched the observer. Trials with non-focal responses but no focal vocalizations were excluded from visual scan and movement analyses but retained for vocal response assessment.

All behavioural data were coded using BORIS v. 8.27.1 (Friard & Gamba, 2016). The temporal window for behavioural analysis was defined based on a prior pilot study and field observations. The analysis period began with the onset of the first playback call and extended five seconds after the end of the second call in the recording. This timeframe was selected to minimize the influence of external disturbances and to focus exclusively on responses directly attributable to the playback stimulus. We extracted the time of occurrence of three key behavioural responses during playbacks: vocal response, walking, and running. A *vocal response* was defined as the emission of at least one contact call by the focal individual during the playback period. Within the broader category of *movement*, we distinguished between walking and *running*. Walking was defined as locomotion involving at least three consecutive steps, with a speed of less than 2 body lengths per second and at least two limbs in contact with the substrate at all times. Running was defined as locomotion at a speed of 2 body lengths per second, including moments when no limbs were in contact with the ground. One body length was operationally defined as the distance from the neck (visually marked by the collar) to the base of the tail.

Two trained observers independently conducted the coding, following a standardised protocol developed in advance. To minimise observer bias, neither observer was aware of the identity of the playback stimulus (i.e., the caller’s sex, rank, or identity) during coding. Inter-observer reliability (IOR) was assessed using a subset of 10 randomly selected videos, independently coded by both observers. Agreement was quantified using Cohen’s Kappa (κ; Cohen, 1960) with the *irr* package in R (Gamer et al., 2019). All behavioural categories yielded acceptable to excellent reliability (κ 0.75), confirming the consistency and robustness of the coding procedure.

### Statistical analysis

We used ordinal Generalized Linear Mixed Models with a cumulative logit link (GLMM; Baayen, 2008; package ordinal, version 2023.12-4.1; Christensen, 2019) to investigate the effects of the absolute dominance rank (*playback rank*) of the animal from which the playback call originated (hereafter “playback (individual)”; Model 1; Supplementary Materials Table S2) and the *rank difference* between the playback individual and the animal to which the playback was presented (hereafter “focal (individual)”; Model 2; Supplementary materials Table S2) on the *movement* response of the focal individual. *Movement* was an ordered categorical response variable indicating whether the focal individual moved during the experimental period, with the possible values “no”, “walk”, and “run” in ascending order. *Rank difference* was calculated by subtracting the *playback rank* from the rank of the focal individual (*focal rank*) for each tested combination. Due to the small number of experiments in which a vocal response occurred (n = 17), we could not meaningfully examine the effect of dominance rank on the occurrence of this response.

In both models, we included *focal sex* and *trial number* as additional fixed effects. In model 1, we also included *focal rank* as a fixed effect to control for its effect. As we reasoned that the effect of rank might depend on *focal sex*, we included the interaction between *playback rank* and *focal sex*. Both models included random intercepts for the *focal identity* and *playback identity*. To avoid overconfident model estimates and maintain the type I error rate at the nominal level of 0.05, we included all theoretically identifiable random slopes (Schielzeth and Forstmeier, 2009; Barr et al., 2013). In model 1, these were the random slopes of *playback rank* and *trial number* within *focal identity*, and *focal rank, focal sex*, and *trial number* within *playback identity*. In model 2, these were the random slopes of *rank difference* and *trial number* within *focal identity*, and *rank difference, focal sex*, and the interaction between *rank difference* and *focal sex* within *playback identity*.

Before fitting the models, we z-transformed *focal rank* and *playback rank* in model 1, *rank difference* in model 2, and *trial number* in both models to a mean of zero and a standard deviation of 1 to facilitate model convergence. Before including *focal sex* as a random slope, we manually dummy-coded and centered it.

In order to avoid cryptic multiple testing (Forstmeier and Schielzeth, 2011) and as an overall test of the effect *play-back rank* (model 1) or *rank difference* (model 2) and its interaction with *focal sex*, we compared each full model (Supplementary Materials, Table S2) with a respective null model which lacked *playback rank* or *rank difference*, respectively, and its interaction with *focal sex* in the fixed effects.

We tested the significance of individual fixed effects terms by dropping them from the model, one at a time, and comparing the fit of the obtained reduced models with that of the respective full model using a likelihood ratio test (R function drop1; Dobson, 2002). We determined 95% confidence limits of model estimates and fitted values by means of parametric bootstraps (N=1000 bootstraps) using a function written by RM. We estimated model stability by dropping each unique focal and playback individual from the data, one at a time, fitting the full model to each of the subsets, and finally comparing the model estimates obtained with those obtained for the full model. This revealed both models to be of good stability). The samples analysed with these models comprised 349 playback experiments made with 18 focals using calls of 21 individuals.

In the results section we show plots with fitted values. These fitted values were determined as weighted means of the response (coding movement category as 0 for no movement, 1 for walking, and 2 for running) whereby the weights were the estimated probabilities for each of the categories to occur. We are aware that these means are not justified given the ordinal nature of the response. However, they might help aiding the interpretation of the model results. For this reason we display them in the plots.

## Results

### Dominance hierarchies

A total of 391 dyadic interactions were tested, of which 96 were tied and excluded from hierarchy construction, leaving 295 decisive interactions for analysis. The dominance network exhibited a high degree of transitivity, with a triangle transitivity proportion (*Pt*) of 0.972 (scaled transitivity index *Ttri* = 0.888), significantly exceeding the expected value for a random network (*Pt* = 0.75; *p* < 0.001, based on 1,000 randomisations), indicating a strong linearity structure in the dominance hierarchy.

Using the I&SI method, we derived a relative dominance rank for each of the 40 individuals (Supplementary materials; Figure S1, Table S1). Aggregated across 10 independent runs, the ranking pattern revealed clear dominance ranks, with adult females consistently occupying the highest ranks (Supplementary Materials, Figure S1, Table S1).

### Playback experiments

We conducted a total of 550 experiments and after applying the exclusion criteria, we were able to label and analyse 378 of them. Out of all experiments analysed, 166 experiments were conducted with females and 212 with males. The IOR test revealed substantial agreement for the labelling of walking (κ = 0.8) and perfect agreement for the labelling of running (κ = 1).

We found a significant effect of *playback rank* on *movement* (full-null model comparison: χ^2^ = 6.11, df = 2, *P* = 0.047).

The interaction between *playback rank* and *focal sex* was also significant (p=0.021; Table 1). In females, *movement* (walking or running) was more common with higher *play-back rank*, whereas in males, no clear trend was observed (Figure 2).

**Table 1.** Model results showing the effect of playback rank on movement in ring-tailed lemurs. Estimates, with their standard errors (SD), 95% confidence limits (CI), significance tests (χ^2^, p), and the range of estimates (min, max) obtained from model stability. *Focal rank, playback rank*, and *trial number* were z-transformed to a mean of zero and a standard deviation (sd) of one. Mean±SD of the original predictors were 18.72 ±and 10.85 (*focal rank*), 16.68± and 9.38 (*playback rank*), and 16.67± and 8.85 (*Trial number*). Sex was dummy coded with female being the reference level.

| Term | Estimate | SE | CI lower | CI upper | $\chi^2$ | df | p | min | max |
| --- | --- | --- | --- | --- | --- | --- | --- | --- | --- |
| 0\|1 | 0.307 | 0.381 | -0.712 | 1.324 | - | - | - | -0.123 | 0.548 |
| 1\|2 | 2.131 | 0.409 | 1.135 | 3.358 | - | - | - | 1.726 | 2.395 |
| Focal rank | -0.096 | 0.311 | -0.945 | 0.743 | 0.095 | 1 | 0.758 | -0.527 | 0.172 |
| Playback rank | 0.401 | 0.175 | 0.092 | 0.800 | - | - | - | 0.349 | 0.460 |
| Sex | -0.590 | 0.632 | -2.298 | 1.024 | - | - | - | -1.510 | -0.193 |
| Trial number | 0.277 | 0.127 | -0.045 | 0.608 | 4.415 | 1 | 0.036 | 0.228 | 0.341 |
| Playback rank×Sex | -0.532 | 0.233 | -1.061 | -0.075 | 5.300 | 1 | 0.021 | -0.611 | -0.464 |

**Fig 2.**
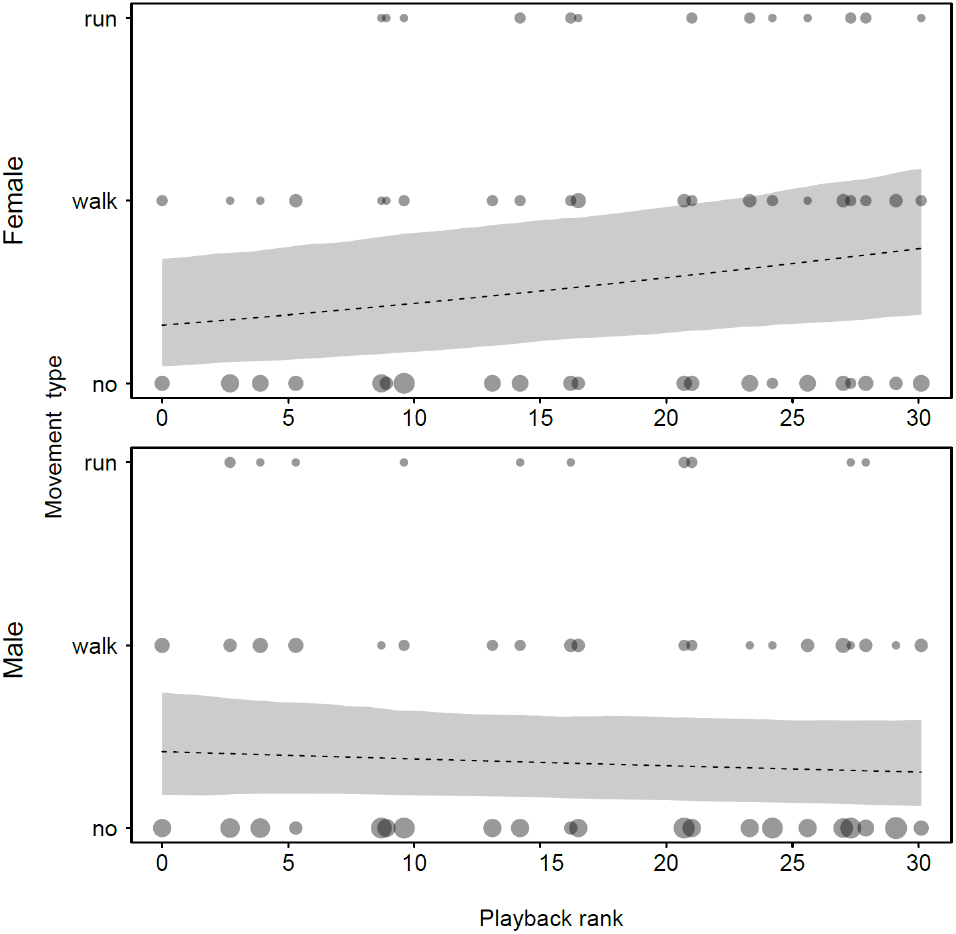
Effects of *playback rank* on the occurrence of *movement* for male and female ring-tailed lemurs. Dashed lines and shaded area show the fitted model and its 95% confidence intervals for all other predictors being in the model being at their average. Dot area is proportional to the number of the respective observations (range 1 to 9).

The effect of *rank difference* on *movement* was marginally non-significant (full-null model comparison: χ^2^ = 5.35, df = 2, *P* = 0.069), suggesting a trend toward an effect of *rank difference*. The interaction between *rank difference* and *focal sex* was significant (β=0.697, p=0.035; Table S3). Plotting the fitted model revealed that in females, *movement* intensity decreased with increasing *rank difference* (i.e., when the playback individual was higher-ranking), while in males, no consistent trend was apparent (Figure S2).

## Discussion

We examined the effects of both caller dominance rank and rank difference between caller and listener on listeners’ behavioural responses to affiliative contact calls in ring-tailed lemurs. We found that females responded more strongly to vocalisations produced by more dominant individuals, initiating movement after hearing contact calls from high-ranking group members. These findings suggest dominant individuals may play an important role in coordinating group movement in this species.

Our results show that adult female ring-tailed lemurs were more likely to move after hearing contact calls from high-ranking callers (Figure 2), suggesting that dominant individuals may play an important role in initiating collective movement. Moan contact calls have previously been suggested to function both in maintaining social relationships and in coordinating group movement in ring-tailed lemurs (Oda, 1996; Bolt and Tennenhouse, 2017). Thus, if high-ranking individuals are more likely to initiate collective movement, responding to their contact calls may be particularly important for females. This interpretation is consistent with findings from other group-living primates. In Alaotran gentle lemurs, higher-ranking individuals were more likely to initiate group movement than lower-ranking individuals (Waeber and Hemelrijk, 2003), while in chacma baboons (*Papio ursinus*), dominant individuals have been shown to occupy consistent leadership roles during group movement (Bracken et al., 2022). Our findings therefore raise the possibility that dominance rank may influence collective decision-making through affiliative vocal communication. However, as the present study did not directly examine naturally occurring group movements, further research is needed to determine whether high-ranking ring-tailed lemurs indeed initiate and lead group movements more frequently than lower-ranking individuals and whether such a role is restricted to females or extends to both sexes.

The effect of caller rank on movement was observed only in females, suggesting that female responses to contact calls may be particularly sensitive to the social hierarchy. In ringtailed lemurs, social relationships among females are closely linked to dominance relationships: spatial proximity and alliances can influence dominance rank and the outcomes of conflicts (Nakamichi et al., 1997). Thus, the importance of female-female social relationships for maintaining positions within the dominance hierarchy may explain why females were particularly responsive to contact calls from high-ranking individuals. Responding to calls from high-ranking females may help maintain proximity to socially important individuals and facilitate the maintenance of social relationships within the female dominance hierarchy.

Females showed a tendency to be more likely to respond to individuals closer to their own position in the hierarchy. Given the highly linear and strict dominance hierarchy characteristic of ring-tailed lemurs) found in our study, responsiveness to individuals of similar rank could potentially contribute to maintaining the stability of this hierarchy. Individuals close in rank may have more directly relevant social relationships, as interactions between them may be particularly important for establishing and maintaining their relative positions (e.g. as partly supported by Snyder-Mackler et al. 2016). Responding to contact calls from such individuals could therefore help maintain appropriate spatial relationships and facilitate the coordination of behaviour among individuals whose positions within the hierarchy are closely connected. This interpretation remains tentative, however, and experimental studies specifically targeting interactions between individuals of different rank distances will be needed to determine whether responses to contact calls contribute to the maintenance of the dominance hierarchy.

In contrast to females, males did not show increased movement responses following calls from higher-ranking individuals. Given that adult males occupy lower positions than all adult females in the ring-tailed lemur hierarchy (Jolly, 1966; Kappeler, 1990), responding to every contact call produced by a higher-ranking female may not provide the same social benefits for males as it does for females. Instead, other aspects of social relationships may be more relevant to male movement decisions. Notably, ring-tailed lemur males have been shown to form particularly strong social bonds with other males during the non-mating season (Gabriel et al., 2014; Gould, 1997), which coincided with the period in which our playback experiments were conducted. It is therefore possible that male responses to contact calls during this period are more strongly influenced by affiliative relationships or bond strength than by dominance rank. In this case, males may be more likely to move towards individuals with whom they share strong social bonds, irrespective of the caller’s position in the dominance hierarchy. Another possibility is that the spatial and social organisation of females during this period reduces the relevance of female-initiated movement for males. Females have been reported to maintain particularly close spatial associations when infants are present (Nakamichi & Koyama, 2000), potentially resulting in female-centred social and movement dynamics that are less closely tied to males. If group movement during our study period was primarily organised around female social relationships, males may consequently have been less motivated to respond to contact calls from high-ranking individuals. Together, these possibilities highlight the importance of considering sex, season, and the broader structure of social relationships when examining the role of dominance in vocal communication. Further studies comparing responses across mating and non-mating seasons and incorporating measures of affiliative bond strength and spatial proximity could help clarify why dominance-related responses were evident in females but not in males.

Some methodological considerations should be taken into account when interpreting our findings. Individuals were more likely to move with increasing trial number, which may indicate an effect of repeated exposure or changes in motivation over the course of the experiment. However, because the presentation order was randomised for each focal individual and was controlled for in our models, this effect is unlikely to have systematically influenced the effects of caller rank or rank difference on movement responses. In addition, as our study relied on playback experiments, the spatial separation between caller and focal individual may have affected behavioural responses and limits the extent to which our findings can be generalised to naturally occurring interactions. Although the absence of a vocal response was unexpected, the rank-related effects on movement indicate that the playback stimuli elicited behaviourally relevant responses. Future studies could therefore examine responses to contact calls in more naturalistic contexts, for example using animal-mounted recording devices to investigate behavioural responses during naturally occurring interactions and group movements.

