## Supplementary Materials for "Effects of caller dominance rank on response to contact calls in ring-tailed lemurs (*Lemur catta*)"

### The effect of dominance rank on behavioural response to playback calls in ring-tailed lemurs (*Lemur catta*)

Table S1: List of all individuals included in the study. Individuals (Focal ID) are sorted by descending dominance rank (inverted averaged I&SI; Focal Rank), sex (f – female, m – male), age (in years) during the study period and indications whether their call was used as playback call (Playback), and whether they were tested in playback experiments (Focal).

| Focal ID | Sex | Age | Focal rank | Playback | Focal |
|---|---|---|---|---|---|
| A | f | 4 | 32.8 | 1 | 1 |
| B | f | 15 | 31.8 | 1 | 1 |
| C | f | 6 | 30.6 | 1 | 1 |
| D | f | 8 | 30 | 1 | 1 |
| E | f | 4 | 29.7 | 1 | 1 |
| F | f | 10 | 28.3 | 1 | 1 |
| G | f | 7 | 26.9 | 1 | 1 |
| H | f | 5 | 26 | 1 | 1 |
| I | m | 10 | 23.7 | 1 | 1 |
| J | m | 8 | 23.4 | 1 | 0 |
| K | m | 12 | 19.2 | 1 | 1 |
| L | m | 7 | 18.9 | 1 | 0 |
| M | m | NA | 16.9 | 1 | 0 |
| N | m | 6 | 15.8 | 1 | 1 |
| O | m | 6 | 12.3 | 1 | 0 |
| P | m | 14 | 11.6 | 1 | 0 |
| Q | m | 4 | 11.4 | 1 | 1 |
| R | m | 18 | 10.2 | 0 | 1 |
| S | m | 4 | 8 | 1 | 1 |
| T | m | 5 | 6.6 | 1 | 1 |
| U | m | 5 | 5.4 | 1 | 1 |
| V | m | 4 | 2.7 | 1 | 1 |
| W | m | 4 | 0 | 0 | 1 |

Table S2. Model formulae for the ordinal generalized linear mixed models used to investigate the effect of dominance rank on movement response of ring-tailed lemurs.

| Model number | Formula |
|---|---|
| Model 1 | movement ~ Focal rank + Playback rank* Focal sex + Trial number + (1+ Playback rank + Trial number \|Focal identity) + (1+ Focal rank + Focal sex + Trial number \|Playback identity) |
| Model 2 | movement ~ Rank difference* Focal sex + Trial number + (1 + Rank difference + Trial number \| Focal identity) + (1+ Rank difference + Focal sex + Rank difference.Focal sex \| Playback identity) |

Table S3: Model results showing the effect of rank difference on movement in ring-tailed lemurs. Indicated are model estimates, with their standard errors (SD), 95% confidence limits (Cl), significance tests (χ2, p), and the range of estimates (min, max) obtained after dropping the levels focal identity and playback identity one at a time. *Rank difference* and *trial number* were z-transformed to a mean of zero and a SD=1. Mean and SD of the original predictors were 2.04 and 14.45 (*rank difference*) and 16.62 and 8.86 (*Trial number*). Sex was dummy coded with female being the reference level.

| Term | Estimate | SE | $CL_{lower}$ | $CL_{upper}$ | ***χ2*** | ***df*** | ***p*** | min | max |
|---|---|---|---|---|---|---|---|---|---|
| Movement 0\|1 | -0.030 | 0.293 | -0.725 | 0.669 | - | - | - | -0.152 | 0.070 |
| Movement 1\|2 | 1.790 | 0.322 | 1.112 | 2.599 | - | - | - | 1.677 | 1.993 |
| Rank difference | -0.584 | 0.263 | -1.247 | -0.094 | - | - | - | -0.661 | -0.511 |
| Sex | -0.785 | 0.379 | -1.763 | 0.063 | - | - | - | -0.897 | -0.626 |
| Trial number | 0.285 | 0.124 | -0.022 | 0.636 | 4.857 | 1 | 0.028 | 0.239 | 0.343 |
| Rank difference:sex | 0.697 | 0.335 | 0.082 | 1.480 | 4.453 | 1 | 0.035 | 0.569 | 0.796 |

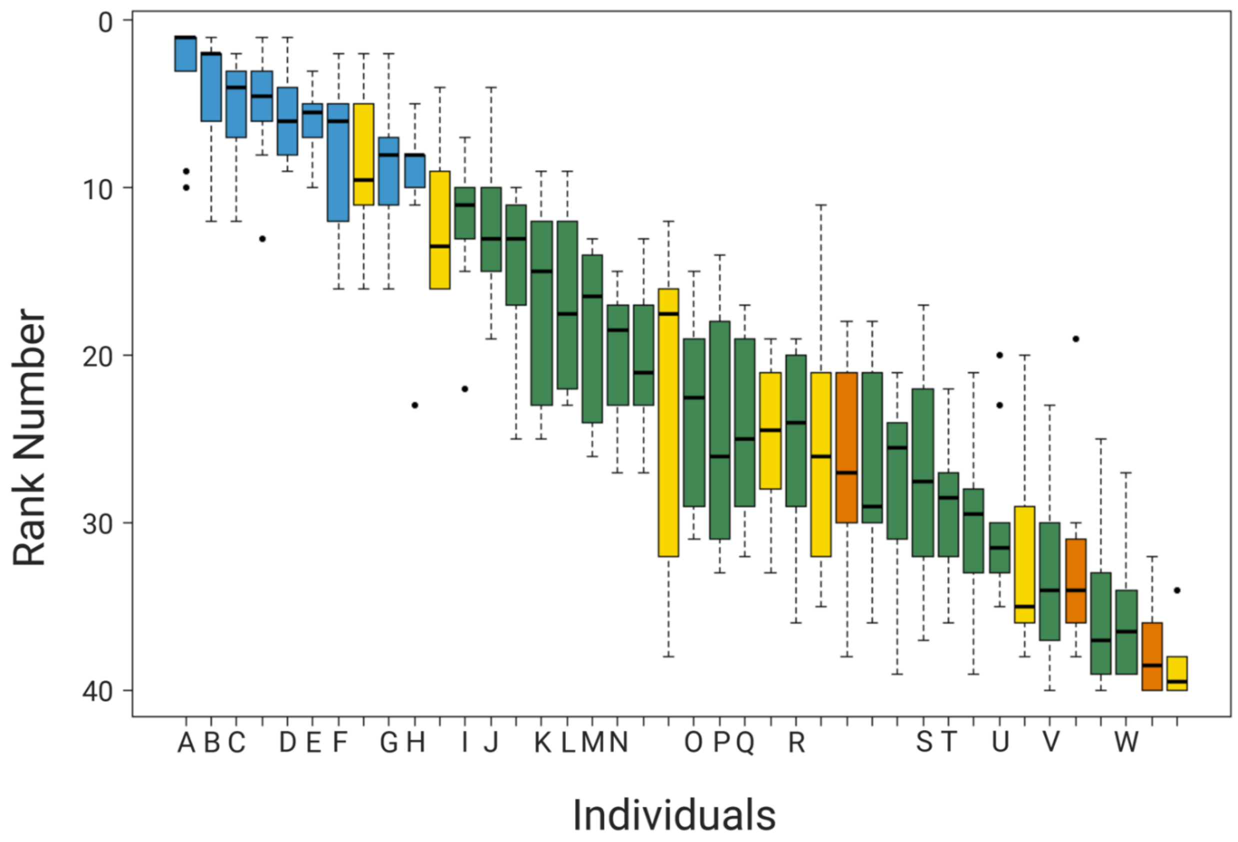


Figure S1. Dominance ranks of ring-tailed lemurs (*Lemur catta*) at Affenwald Straussberg. Individual ranks across 10 independent I&SI hierarchy iterations are plotted, ordered by average I&SI rank. Colors denote age and sex classes: adult females (blue), adult males (green), juvenile females (yellow), and juvenile males (orange). The vertical spread reflects rank variability across iterations. Letters correspond to focal individuals in the playback experiments (Table S1).

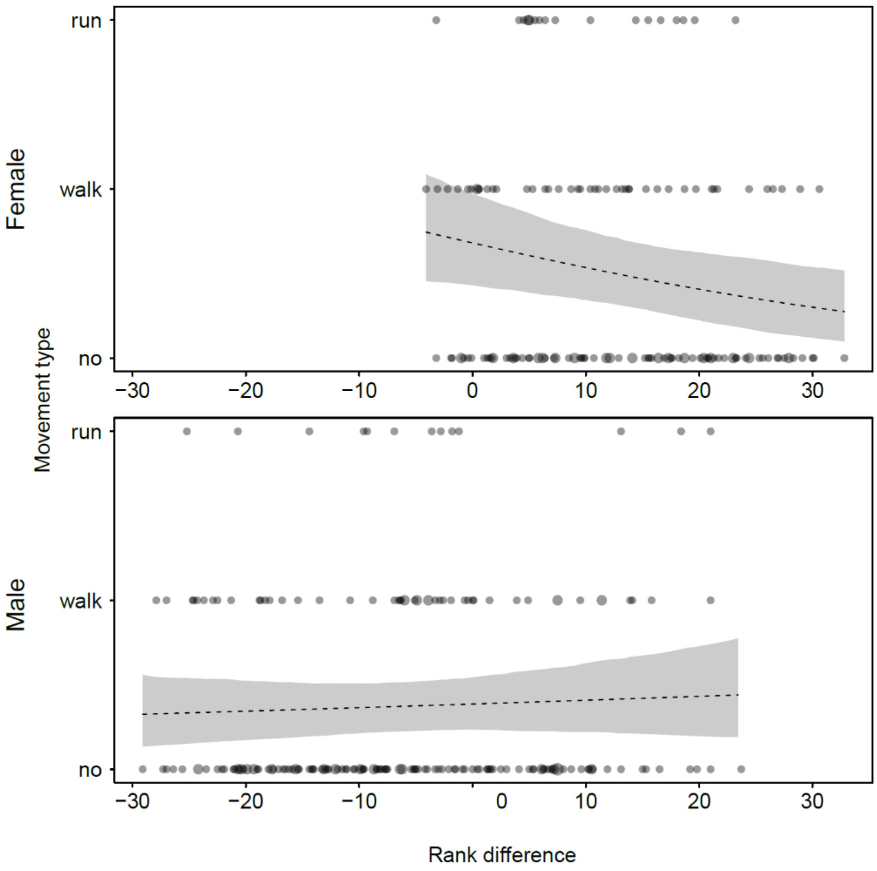


Figure S2: Effects of *rank difference* on the occurrence of *movement* for male and female ring-tailed lemurs. Dashed lines and shaded area show the fitted model and its 95% confidence intervals for all other predictors being in the model being at their average. Dot area is proportional to the number of the respective observations (range 1 to 3). The fitted models were only plotted across observed rank differences.
